# Metastable regimes and tipping points of biochemical networks with potential applications in precision medicine

**DOI:** 10.1101/466714

**Authors:** Satya Swarup Samal, Jeyashree Krishnan, Ali Hadizadeh Esfahani, Christoph Lüders, Andreas Weber, Ovidiu Radulescu

## Abstract

The concept of attractor of dynamic biochemical networks has been used to explain cell types and cell alterations in health and disease. We have recently proposed an extension of the notion of attractor to take into account metastable regimes, defined as long lived dynamical states of the network. These regimes correspond to slow dynamics on low dimensional invariant manifolds of the biochemical networks. Methods based on tropical geometry allow to compute the metastable regimes and represent them as polyhedra in the space of logarithms of the species concentrations. We are looking for sensitive parameters and tipping points of the networks by analyzing how these polyhedra depend on the model parameters. Using the coupled MAPK and PI3K/Akt signaling networks as an example, we test the idea that large changes of the metastable states can be associated to cancer disease specific alterations of the network. In particular, we show that for model parameters representing protein concentrations, the protein differential level between tumors of different types is reasonably reflected in the sensitivity scores, with sensitive parameters corresponding to differential proteins.

## 1 Introduction

Precision medicine is an emerging concept in healthcare that aims to adapt the therapy to patient specificity [10]. The need for this paradigm change is justified by the strong differences among patients suffering of seemingly the same disease which leads to strong variability of the response to treatment. The implementation of such a strategy relies on the development of new tools that allow to understand the origin of the inter-individual differences and to predict the individual response to a particular treatment strategy.

It is now largely considered that at least part of the inter-individual differences are genetic and therefore can be detected by genome sequencing. However, although differences in a DNA sequence, such as single nucleotide polymorphism and copy number variations, are relatively easy to detect, their consequences are very difficult to predict [30, 35, 11]. Mathematical models for cell physiology such as protein interaction networks and chemical reaction networks (CRN) can be used to understand the impact on the phenotype of a change in the protein function or expression level. Static protein interaction networks were used to understand differences between patients by mapping the genotype differences onto the network and identifying significantly enriched modules [8]. This method, based on the hypothesis of modularity of the biological function, has its limitations. Most importantly, it is unable to predict differences in the phenotype produced by alterations of the same pathway.

In oncology, the concentration of alterations in the same pathway is not uncommon, especially for hub signaling pathways such as MAPK and PI3K/Akt [34]. These pathways are deregulated in more than 60% of all cancers; these deregulations imply multiple changes involving several proteins. Pathway redundancy and multiple feed-back regulation are obstacles against cancer targeted therapies; inhibition of one oncogene can trigger compensatory effects elsewhere [19, 13]. Furthermore, deregulation of signaling is a dynamical phenomenon, affecting the time dependent response of signaling proteins to stimuli. Therefore, in order to predict the effect of perturbations on such complex systems, one needs quantitative dynamical models. Signaling pathway dynamics can be conveniently modeled by ordinary differential equations (ODEs) resulting from the chemical kinetics of CRNs.

In the context of CRN dynamics, differences between patients or between tumors can be modeled as differences of parameter values. It is reasonable to think that only the differences of the sensitive parameters, i.e., those parameters whose change induces significant effect, matter for the differences in phenotype. Sensitivity analysis is a general method allowing to detect sensitive parameters and the capacity of the model to resist changes [37]. Sensitivity analysis can be applied to steady states or to computation tree logic (CTL) formulas or ad hoc descriptions of properties of time dependent protein concentrations such as oscillations, peaks, etc. [28, 36, 18]. However, in general it is not clear which dynamical property is important for the biological function. Therefore, it is interesting to look for global methods allowing to test the effect of parameter changes on all the features of the dynamics.

A possible global description of a dynamical system can be the set of its point attractors (stable steady states). In the context of boolean network models (an alternative to ODE models) point attractors have been used to characterize cell types and changes of their number were interpreted as cell fate decisions [15]. However, the set of point attractors is difficult to compute for large networks. Moreover, the knowledge of point attractors is not sufficient for reconstructing the transient propagation of a signal through the network.

We have recently shown that biochemical networks in general, and signaling networks in particular, have also metastable regimes, defined as regions of the phase space where the system is slow and spends a long time [33]. A possible trajectory passes from one metastable regime to another, either stopping in a steady state or visiting periodically one or several slow metastable regimes along a limit cycle attractor. Thus, the set of metastable regimes and the possible transitions between them provides a richer picture of the qualitative dynamics of the network [33, 26]. We also showed how to use tropical geometry methods in order to compute, without simulating trajectories, all the metastable regimes of a given model with polynomial ODE dynamics [32, 33, 26]. In [33] we showed that data from aggressive and non-aggressive tumors correspond to different metastable regimes of the TGF-*β* signaling network.

In this paper, we provide a new application of tropical geometry methods. In computational models of signaling pathways, some proteins represent model variables and other proteins represent model parameters. Of course, not all model parameters can be associated to protein levels, many of them are kinetic parameters. In [33] we focused on those proteins that are model variables and directly compared their levels in metastable regimes to data. Here, we use tropical geometry to detect sensitive parameters that have significant global effect on the network dynamics. For those parameters that correspond to proteins we compare the sensitivity scores to differential protein levels from data. Changing sensitive parameters can bring the network to tipping points where the network behavior changes drastically. In order to test this idea in a biomedical framework we check if these sensitive parameters are significantly changed in cancer disease.

The structure of this paper is the following. The second section recalls the mathematical formalism of tropical geometry and introduces the branches of tropical equilibration that are proxies for metastable regimes. The third section is dedicated to the methods used for computing tropical equilibration branches and for performing sensitivity analysis. The fourth section presents the results of this analysis on a MAPK and PI3K/Akt model from the literature and the comparison of sensitive parameters with differential proteome data from different cancers. The last section is a discussion of the results.

## 2 Theory: Tropical equilibrations of chemical reactions networks with rational rate functions

In this section we introduce the main concepts relating to geometry and dynamics.

We consider chemical reaction networks described by rational kinetic laws (the reaction rates are fractions whose denominator and numerator are polynomials in the species concentrations). Examples of such kinetic laws include, but are not restricted to, mass action law, Michaelis-Menten law and its various generalizations, Hill law with integer Hill index.

After computing a common denominator of rates of reactions acting on the same species, the CRN kinetics are described by a system of differential equations:

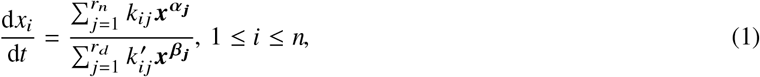

where *k_ij_*, 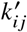 are products of kinetic constants and integer stoichiometric coefficients, *x*_i_ > 0 are variable concentrations, *α_j_* = (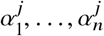), *β_j_* = (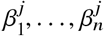) are multi-indices, and ***x^α_j_^*** = 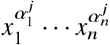 ***x^β_j_^*** = 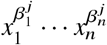.

In general, the system (1) has a number of semi-positive conservation laws, i.e., linear combinations of variables that are constant on any trajectory:

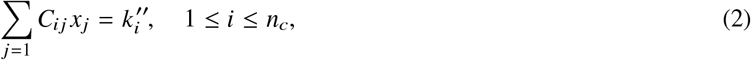

where *C*_ij_ ≥ 0, 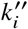 > 0.

We replace now the exact values of parameters by their orders of magnitude which are supposed to be known. Usually, orders of magnitude are approximations of the parameters by integer powers of ten and serve for rough comparisons. Our definition of orders of magnitude is based on the equations *k*_ij_ = *k_ij_ε^γij^*, 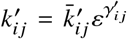, 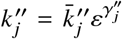, where 0 < ε < 1. The exponents *γ,γ^ʹ^, γ^ʺ^* are considered to be integer or rational. For instance, the approximation

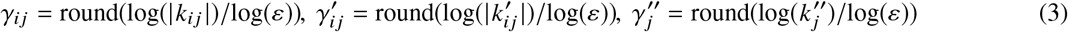

produces integer exponents, whereas *γ_ij_* = round(*d* log(~*k_ij_*)~/log(*ε*))/*d*, 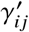 = round(*d* log(~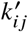~/log(*ε*))/*d*, 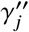 = round(*d* log(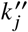)/log(*ε*))/*d* produces rational exponents, where “round” stands for the closest integer (with half-integers rounded to even numbers) and *d* is a strictly positive integer. When *ε* = 1/10, our definition provides the usual decimal orders.

In this study, orders of magnitude of the kinetic parameters are supposed to be known. In contrast, species orders vary in time and have to be computed. To this aim, the species concentrations are first represented by orders of magnitude defined as

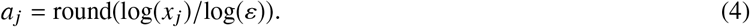

More precisely, one has *x_j_* = *x_j_ε^a_j_^*, where *x_j_* has zero order (unity). The definition (4) restricts species orders to integers; however, rational orders are also acceptable. Because log(*ε*) < 0, (4) means that species orders and concentrations are anti-correlated (large orders mean small concentrations and vice versa).

Then, network dynamics are described by the rescaled ODE system

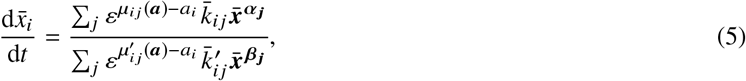

where *µ_ij_*(***a***) =*σ_ij_* + 〈***a*, *α_j_***〉, 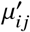(*a*) = 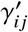 + 〈***a*, *β_j_***〉 and 〈·, ·〉 stands for the dot product.

The rescaled conservation laws read

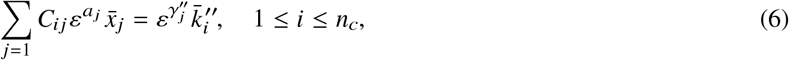

where we considered that the *C_ij_* have zero order.

The numerator and denominator of the r.h.s. of each equation in (5) are sums of multivariate monomials in the concentrations. The orders *µ_ij_* indicate how large these monomials are in absolute value. A monomial of order *µ_ij_* dominates another monomial of order *µ_ij_*ʹ if *µ_ij_* < *µ_ij_ʹ*.

There is always at least one dominant monomial for the numerator and one for the denominator. We are interested in the situation when the denominator has at least two dominant monomials, one positive and the other negative. This situation implies compensation of large forces on variable *i* and as shown elsewhere [33], leads to a good proxy for metastable states. Furthermore, the orders of magnitude are constrained by the conservation laws. This justifies the following definition:

*The tropical equilibration problem* consists of the equality of the orders of at least two monomials, one positive and another negative, in the numerator of the differential equations of each species and in each semi-positive linear conservation law. This condition allows us to compute the concentration orders defined by (4). More precisely, we want to find a vector **a** such that min

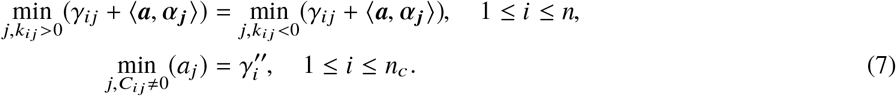

Eq. (7) is related to the notion of a *tropical hypersurface*. A *tropical hypersurface* is the set of vectors a ∈ ℝ^n^, such that the minimum min_j_(*γ_ij_* + 〈***a*, *α_j_***〉) is attained for at least two different indices *j* (with no sign conditions). *Tropical prevarieties* are finite intersections of tropical hypersurfaces. Therefore, our tropical equilibrations are subsets of tropical prevarieties.

In order to find the solutions of the system (7) we can explore combinatorially trees of solutions resulting from various choices of minimal terms and write down inequalities for each situation. Because a set of inequalities defines a polyhedron, the set of tropical equilibration solutions forms a set of polyhedra in ℝ^n^.

To set these ideas down let us use a simple chemical network example, the Goldbeter-Koshland kinetics:

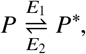

where *P*, *P*\*, *E*_1_, *E*_2_ represent a protein, its modified form, and two enzymes, respectively.

The two enzymatic reactions of the model have Michaelis-Menten kinetics and it results

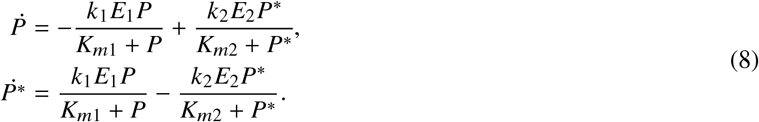

where *k*_1_, *k*_2_, *K*_m1_, *K*_m2_ are positive kinetic constants.

The system (8) has a semi-positive linear conservation law

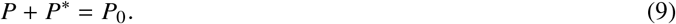

Orders of variables and parameters are as follows: *P* = *P*ε*^a_1_^*, *P*\* = *P*\* ε*^a_2_^*, *k*_1_*E*_1_ = *v*_1_ *ε^γv^*^1^, *k*_2_*E*_2_ = *v*_2_*ε^γ_v_2^*, *K*_m1_ = *K*_m1_*ε^γ_m_1^*, *K*_m2_ =*K*_m2_*ε^γ_m_2^*, *P*_0_ = *P*_0_*ε^γ_p_0^*.

After finding the common denominator of the fractions in (8) we get

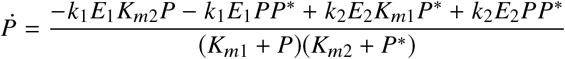

and the tropical equilibration equations for the Goldbeter-Koshland example read

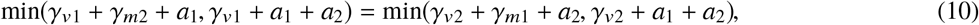

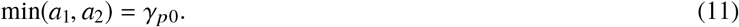

By examining all the cases (10) and (11) we find values for *a*_1_ and *a*_2_, but also conditions that the parameters must satisfy for these solutions. We list below all the structurally stable solutions (solutions whose existence does not depend on equalities among parameter orders):

1. *a*_1_ = *γ_p0_*, *a*_2_ = *γ_p0_* + *γ_v1_* − *γ_v2_* + *γ_J2_* − *γ_J1_* under the conditions *γ*_*J*1_ ≥ 0 and *γ*_v1_ − *γ*_v2_ ≥ max(*γ_J1_*, *γ_J1_* − *γ_J2_*),
2. *a*_2_ = *γ_p0_*, *a*_1_ = *γ_p0_* + *γ_v2_* − *γ_v1_* + *γ_J1_* − *γ_J2_* under the conditions *γ*_*J*2_ ≥ 0 and *γ*_v2_ − *γ*_v1_ ≥ max(*γ_J1_*, *γ_J1_* − *γ_J2_*),
3. *a*_1_ = *γ_p0_*, *a*_2_ = *γ_p0_* + *γ_v1_* − *γ_v2_* + *γ_J2_* under the conditions *γ_J1_* ≤ 0 and *γ_v1_* − *γ_v2_* ≥ max(−*γ_J2_*, 0),
4. *a*_2_ = *γ_p0_*, *a*_1_ = *γ_p0_* + *γ_v2_* − *γ_v1_* + *γ_J1_* under the conditions *γ_J2_* ≤ 0 and *γ_v2_* − *γ_v1_* ≥ max(−*γ_J1_*, 0),

where *γ_j1_* = *γ_m1_* − *γ_p0_*, *γ_j2_* = *γ_m2_* − *γ_p0_*. For this model, for the given parameter values, there is only one tropical equilibration solution that follows closely the unique stable state state of the model when the model parameters are perturbed (see Fig. 1). Interestingly, the well-known zero-order ultrasensitivity [14] of this model, occurring when both enzymes are saturated (i.e., when *γ_j1_* ≤ 0, *γ_j2_* ≤ 0), corresponds to a jump discontinuity of the tropical solution (see Fig. 1).

**Fig. 1:**
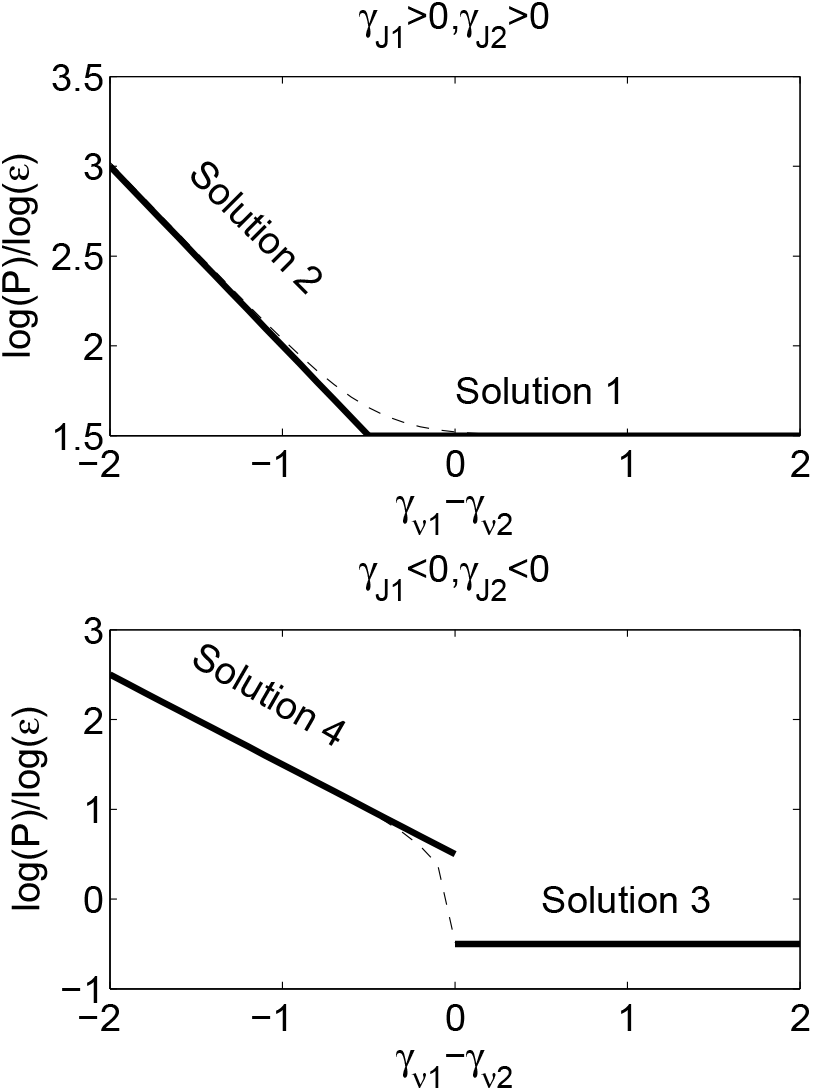
Tropical equilibration solutions for the Goldbeter-Koshland (GK) model. This simple model has one stable steady state. The tropical equilibration solutions (thick lines) follow closely the positions of the stable steady state (dashed line) when the parameter *k*_1_*E*_1_/*k*_2_*E*_2_ changes. The two situations *γ*_*J*_1__≥ 0, *γ*_*J*_2__≥ 0 and *γ_J_1__* ≥0, *γ_J_2__* 0 correspond to the well known [14] unsaturated and saturated (ultrasensitive) regimes of the GK model, respectively.

*Branches of tropical equilibrations*. For each equation *i*, let us define

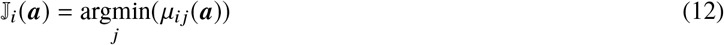

in other words J_i_ denotes the set of indices *j* of monomials having the same minimal order for a given *i*.

We say that two tropical equilibrations ***a*_1_, *a*_2_** are equivalent iff 𝕁_i_(***a*_1_**) = 𝕁*_i_*(***a*_2_**), for all *i*. Equivalence classes of tropical equilibrations are called *branches*. A branch *B* with index sets 𝕁_i_ is *minimal* if, for any branch ***B*^ʹ^** of index sets 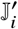, the relation 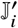 ⊂ 𝕁_i_ for all *i* implies ***B***^ʹ^ = ***B*** or ***B*^ʹ^** = **0̸**. Closures of equilibration branches are defined by a finite set of linear inequalities, which means that they are polyhedral complexes. Minimal branches correspond to maximal dimensional faces of the polyhedral complex. The incidence relations between the maximal dimensional faces (*n*−1 dimensional faces, where *n* is the number of variables) of the polyhedral complex define the *connectivity graph*. More precisely, minimal branches are the vertices of this graph. Two minimal branches are connected if the corresponding faces of the polyhedral complex share an *n* − 2 dimensional face. In terms of index sets, two minimal branches with index sets 𝕁_i_ and 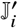 are connected if there are index sets 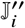 of an existing non-minimal branch such that 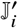 ⊂ 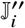 and 𝕁_i_ ⊂ 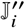 for all *i*. A zero dimensional minimal branch is a point, like in the case of the Goldbeter-Koshland model when it corresponds to a steady state of the model. A non-zero dimensional minimal branch usually corresponds to a slow invariant attractive curve or surface (sufficient conditions for such a situation can be found in [27]). Examples of models with non-zero dimensional branches corresponding to attractive invariant manifolds can be found in [24, 32, 27, 33]. Slow attractive invariant manifolds are metastable regimes because the dynamical system defined by Eq. (1) spends considerable time on them before eventually leaving them for another invariant manifold.

## 3 Methods

### 3.1 Computation of minimal branches

For a given model and set of model parameters, the minimal branches were computed by the algorithm described in [31, 33]. The tropical solution set is the union of polyhedra where the polyhedra represent minimal branches.

### 3.2 Generation of perturbed parameter orders

For a given model, the nominal tropical solution set *M* was computed with the given model parameters. Thereafter, the perturbed tropical solution sets 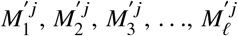 were computed by varying the orders of the given parameter *k*_j_. Here, *ℓ* = 6 and the perturbed parameter orders are *γ_j_* −3, *γ_j_* −2, *γ_j_* −1, *γ_j_* +1, *γ_j_* +2, *γ_j_* +3, respectively. The perturbation of multiple parameter orders in such a framework is straightforward.

### 3.3 Identification of parameter sensitivity scores

#### 3.3.1 Computation of distances

For distance computation, we applied the following constraints to the concentration orders *a_j_* (cf. (4)), thereby defining a feasible region Γ within the polyhedra for sampling purposes:

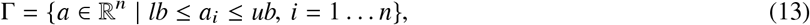

where *lb* and *ub* refers to upper and lower bounds, respectively. Finally, the distances between 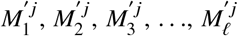 were computed and are represented by the vector *D^j^* of dimension *ℓ* as defined in the following manner

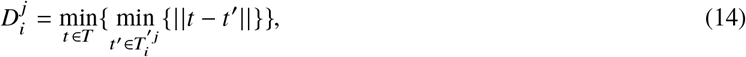

where *T* and 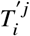 denote the sets of representative point(s) sampled from the polyhedra in *M* ∩ Γ and 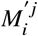 ∩ Γ, respectively.~~ · ~~ is the *L_p_* norm distance metric. The philentropy R-package with Minkowski distance method [12] was used to implement it. We tested our findings with different values of the *L_p_* norm (as suggested in [7]). The idea of such a distance computation technique was motivated by collision detection approach in motion planning (cf. Chapter 5 in [20]). For a given *M*, *T* was computed as follows:

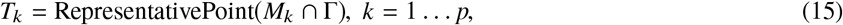

where *p* is the number of polyhedra in the solution set *M*. RepresentativePoint(·) is a function that generates random sample point(s) satisfying the inequality conditions for the given polyhedron and computes the component-wise mean. In case of non-zero dimensional polyhedra, we sampled 3000 random samples points from an infinite number of feasible points.

If instead of ℝ^n^ distances one is interested in the effect of perturbations on a particular target variable *X_m_*, then the sets *T_k_* should be replaced by their projections on the corresponding axis. More precisely, instead of the concentration order vector **a** = (*a*_1_, … , *a_n_*) ∈ *T_k_*, consider the scalar *a_m_* representing the concentration order of the species *X_m_*. We will denote these projected distances as 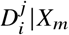

#### 3.3.2 Tropical sensitivity score

We denote the mean

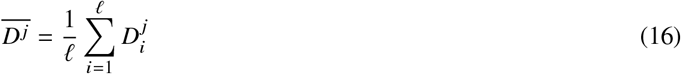

as the parameter sensitivity score for *k_j_*. Similarly, we define

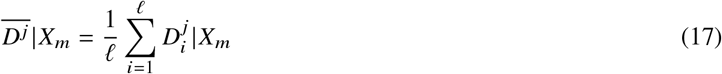

as the parameter sensitivity score for *k_j_* in the direction of the variable *X_m_*. A higher score signifies a more sensitive parameter in a relative scale. However, sensitivity scores can not be compared between (17) and (16).

#### 3.3.3 Availability

TROSS (Tropical Sensitivity Scores), the software to compute the sensitivity scores is available at https://github.com/JRC-COMBINE/TROSS. It uses PtCut to compute the tropical solutions [23].

## 4 Results

In this section, we test the idea that large changes of the metastable states can be associated with cancer disease specific alterations of the network. We selected a biochemical reaction network known to be involved in cancer and computed the tropical minimal branches with methods exposed in the Sections 2 and 3.1, thereby approximating its metastable states. Large changes in tropical solutions are achieved by changing the parameter values from a given set of nominal values and quantifying the resulting change through normalized sensitivity scores. The comparison between these scores and differential proteomics data obtained from different cancer conditions is presented.

More precisely, we perform a diagnostic test to check if the protein levels whose variations between different cancer conditions are large are also strongly sensitive parameters of the model and that proteins with small variation are low sensitivity parameters. The accuracy of such a test is based on the receiver operating characteristic (ROC) curve and the area under the ROC curve (AUC) measures if the ordering in the parameter sensitivity scores (sensitive/not sensitive) is actually preserved in the differential expression of proteins (high/low) in the data.

### 4.1 Biochemical reaction network

The biochemical model was obtained from the Biomodels database [21] with identifier BIOMD0000000146. The model, introduced in [16], was motivated by experimental works on the Heregulin stimulated ErbB receptor and demonstrates the Akt-induced inhibition of the MAPK pathway via phosphorylation of Raf-1. We have chosen this model because it is one of the few models in the Biomodels database showing crosstalk of two pathways important in cancer. A very similar model was used in [19] to investigate the possibilities of targeted combination therapy.

This CRN model has 33 species and 34 reactions: 21 reactions have Michaelis-Menten kinetics and 12 have mass action kinetic laws. The chemical kinetics is composed of 33 ODEs with rational r.h.s. terms. Using MATLAB Symbiology we computed 11 semi-positive conservation laws. Using the conservation laws and the denominator of common denominator expression for each ODE we generated a system of 44 polynomials for which we computed tropical equilibrations.

The CRN has 78 parameters to which we added 11 more parameters corresponding to the initial values of the conservation laws, computed from the initial concentrations of the species.

Among the 89 parameters not all of them amount to expression level of genes and proteins. The parameters that can be directly associated to genes and proteins were chosen according to the following simple method:

- All the concentration laws have a meaning as total amount of kinases and were associated to the corresponding protein kinase: *k*_79_–*k*_89_. Exception is made by *k*_80_ that represents conservation of the complex AKT-PI-P and can not be directly associated to a protein.
- Some other parameters belong to dephosphorylation reactions and are proportional to constant phosphatase concentrations: *k*_77_, *k*_78_. These parameters were associated to the corresponding protein phosphatase.

In the Sections 4.3.3 and 4.4.2 the sensitivity scores of the chosen parameters are compared to differential levels of the corresponding proteins found in databases. To this aim we associate to each parameter the protein whose level is represented by the value of the parameter. In models, variable and parameter names often correspond to protein families. For instance, the parameter *k*_77_ coding for a MAPK phosphatase, generically designates the DUSP family. Thus, the mapping from parameters to proteins is one-to-many.

### 4.2 Computation of distances

For the given (default) parameter values of this model, we obtained two minimal branches, each of which were of dimension 2. The existence of non-zero dimensional branches suggests that all of them are metastable regimes (cf. Section 2). The minimal branches along with the parameter perturbations were performed as per the steps described in Section 3 for *ε* = 1/11. For sampling, we fixed the *ub* = round(log(10^−20^) log(*ε*)) and *lb* = round (log(10^10^)/log(*ε*)), so that it covers a broad range of concentration orders applicable in different application scenarios. Furthermore, we normalized the parameter sensitivity scores between 0 and 1 in the following manner

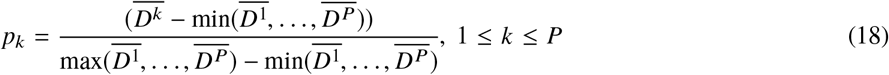

where *D^k^* (cf. Eq. (16), (17)) and *pk* are the parameter sensitivity and normalized parameter sensitivity score of *k*-th parameter respectively. *P* = 89 (total number of model parameters). The histogram depicting the normalized parameter sensitivity scores is shown in Fig. 2 for the Euclidean distance, i.e., using the *L_p_* norm with *p* = 2. For this distance, the distribution of sensitivity scores is rather uniform and most sensitive parameters can not be sharply separated from the others. We tested the robustness of the sensitivity scores by varying the *L_p_* norm (cf. Supplementary Fig. 3). One can notice that a small value of the *L_p_* norm preserves the order and sharpens the differences between sensitivity scores, emphasizing a few strong sensitivity parameters. The normalized sensitivity scores for various model parameters are presented in Fig. 2 for three distances *D_1_* computed with Eq. (16), and *D_2_*, *D_3_* computed with Eq. (17), for *L_p_* norm with *p* = 2 (cf. Supplementary Table 3 for other values of the *L_p_* norm). The distances *D_2_* and *D_3_* are in projection on a 1D axis, therefore do not depend on the value of the *L_p_* norm, except for minor variations due to the random sampling of the polyhedra.

**Fig. 2:**
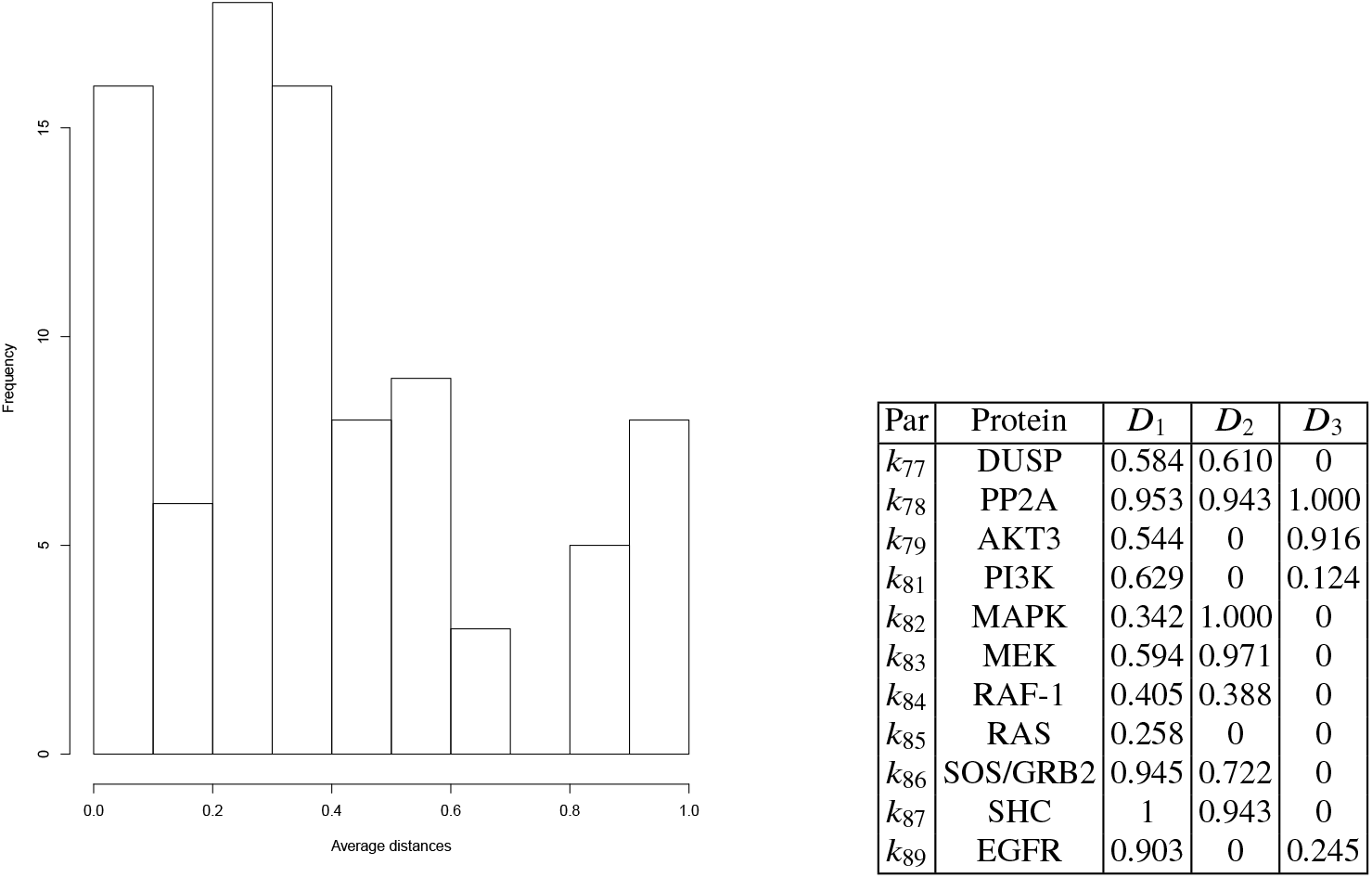
Histogram (in the left) depicts the distribution of normalized average distances (sensitivity scores) of 89 model parameters obtained from BIOMD0000000146 and the *L_p_* norm with *p* = 2 (cf. Section 3.3.1) and distance version *D*_1_. Table (in the right) lists the parameters in BIOMD0000000146 that can be mapped to proteins. Compared to the Biomodels version, the parameters were renumbered from 1 to 89, including the 11 conservation laws. The parameter sensitivities are computed with the *L_p_* norm with *p* = 2 and are provided as normalized average distances in 3 versions: *D*_1_ full distance, *D*_2_ distance along MAPK-PP axis, *D*_3_ distance along AKT-PI-PP axis. *D*_1_ is computed using Eq. (16), whereas *D*_2_ and *D*_3_ are computed based on Eq. (17) with MAPK-PP and AKT-PI-PP as target variables, respectively.

**Fig. 3:**
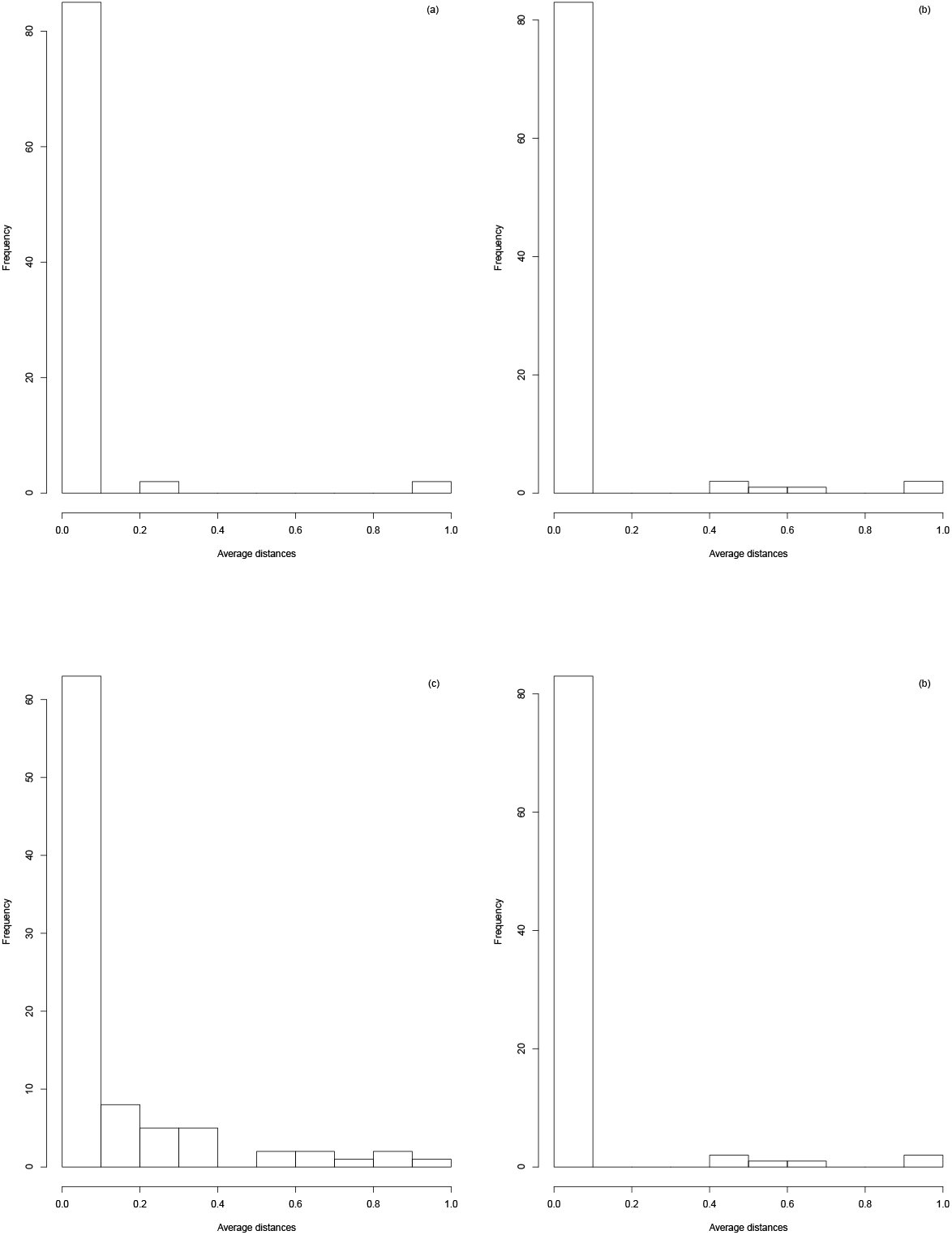
Histograms depict the distribution of sensitivity scores of 89 model parameters obtained from BIOMD0000000146. The histograms (a), (b), (c), (d) correspond to *L_p_* norm values of 0.01, 0.1, 0.5 and 1 respectively (cf. Section 3.3.1 in main text).

### 4.3 TCPA Proteomics Data

We focused on the tissue samples pertaining to The Cancer Genome Atlas (TCGA). TCGA is among the biggest collaborations with the aim of genomic research in cancer and comprises of more than two petabytes of genomic data [5], which makes it an important benchmarking dataset in the bioinformatics community. Since in this work we aimed to study the protein differential levels, we turn to The Cancer Proteome Atlas (TCPA), which contains 8167 tumor samples in total, mainly consisting of TCGA tumor tissue sample sets [6].

#### 4.3.1 Data pre-processing

We selected L4 (Replicates-based normalization) data files from [22] with at least two groups of patient cohorts. This resulted in four cancer datasets, namely, Breast invasive carcinoma (BRCA), Pheochromocytoma and Paraganglioma (PCPG), Skin Cutaneous Melanoma (SKCM) and Thyroid carcinoma (THCA), respectively. BRCA dataset consisted of primary tumor, metastatic tumor and normal samples whereas the rest comprised of primary and metastatic samples. The samples are relevant to the TCGA Research Network.

#### 4.3.2 TCPA Differential Protein Expression Analysis

Out of 89 model parameters, we could associate 9 of them (see Fig. 2) with the datasets where more than one protein may be mapped to the same parameter. Thereafter, we extracted the data for the 17 proteins associated with these 9 parameters and performed a Wilcoxon rank sum test [17] between the primary versus normal samples in BRCA and primary versus metastatic samples for other datasets. The P-values were adjusted based on Benjamini and Hochberg correction method [9]. P-values smaller than a threshold indicate significant differential protein expression. Only BRCA and SKCM datasets resulted in significant proteins, where significance is defined at various P-value thresholds, namely, 5e-05, 5e-04, 0.005, 0.05. In case several proteins are mapped to a single model parameter, the parameter is considered to be significant if it is mapped to at least one of the significant proteins. The adjusted P-values are reported in Table 1.

**Table 1:** Adjusted P-values computed among various subtypes of breast cancer from CPTAC database (Basal-like, HER2-enriched, Luminal-B, Luminal-A) and BRCA (normal versus primary samples), SKCM (metastatic versus primary samples) cancers from TCPA database (cf. Section 4.4.1, 4.3.2 for details). In case of multiple proteins mapping to the same parameter, the one with the lowest P-value is reported.

| Par | Protein | P-values<br>Basal-like<br>vs<br>HER2-enriched | P-values<br>Basal-like<br>vs<br>Luminal-B | P-values<br>HER2-enriched<br>vs<br>Luminal-B | P-values<br>Basal-like<br>vs<br>Luminal-A | P-values<br>BRCA | P-values<br>SKCM |
| --- | --- | --- | --- | --- | --- | --- | --- |
| k77 | DUSP | 0.772 | 0.521 | 0.946 | 0.994 | 0.355 | 0.178 |
| k78 | PP2A | 0.487 | 0.160 | 0.909 | 0.028 | 0.004 | 0.068 |
| k79 | AKT3 | 0.400 | 0.061 | 0.937 | 0.892 | 0.037 | 0.396 |
| k81 | PI3K | 0.321 | 0.011 | 0.909 | 0.017 | 0.006 | 0.0121 |
| k82 | MAPK | 0.557 | 0.227 | 0.909 | 0.028 | 0.006 | 0.0121 |
| k83 | MEK | 0.487 | 0.227 | 0.909 | 0.110 | 1.852e-06 | 0.068 |
| k84 | RAF-1 | 0.557 | 0.227 | 0.909 | 0.717 | 6.84e-09 | 0.012 |
| k85 | RAS | 0.487 | 0.106 | 0.909 | 0.308 | 6.848e-09 | 0.696 |
| k86 | SOS/GRB2 | 0.006 | 0.017 | 0.909 | 0.018 | 1.092e-06 | 0.702 |
| k87 | SHC | 0.487 | 0.467 | 0.909 | 0.965 | 1.092e-06 | 0.702 |
| k89 | EGFR | 0.487 | 2.000e-05 | 0.018 | 0.002 | 2.306e-07 | 0.068 |

#### 4.3.3 Validating parameter sensitivity scores with TCPA database

We compared model parameter sensitivity scores and differential protein expression. To this aim, we defined a diagnostic test to assess the ability of our method to correctly identify sensitive/insensitive parameters with respect to their differential expression in the proteomics data. We treated the normalized parameter sensitivity scores (cf. Section 4.2) as predicted class probabilities ranging from 0 to 1. The parameters with values close to 1 are more sensitive than the ones close to 0. Finally, we perform a diagnostic test to quantify the extent to which the predicted class probabilities overlap with the significantly (actual class label of 1) and non-significantly (actual class label of 0) expressed proteins in our datasets. The test is performed by computing the AUC of the ROC curve using R [29]. The major benefit of using AUC is that the method does not require cut points in order to define predicted class labels from predicted class probabilities [25]. An AUC above 0.5 means that the method performs better than random guessing. As the definition of significantly expressed proteins, as well as the *L_p_* norm values, are arbitrary, i.e., depends on a pre-defined threshold, we investigated the changes in AUC values by varying the P-value thresholds while keeping the *L_p_* value constant and vice versa. The findings are reported in Table 2 and Supplementary Tables 4, 5. The distribution of AUC values are shown in Supplementary Fig. 4, 5.

**Table 2:** Left table: In each column, the median AUC values are reported by averaging over different *L_p_* norm values (*p* = 0.01, 0.1, 0.5, 1, 2) while keeping the P-value threshold at 5e-02. SD represents the standard deviation. AUC values at NA entries could not be computed due to the absence of significant proteins at the particular P-value threshold. Right table: In each column, the median AUC values are reported by averaging over different P-value thresholds (5e-05, 5e-04, 0.005, 0.05) while keeping the *L_p_* norm at *p* = 2. SD values at NA entries could not be computed due to the presence of significant proteins only at a single P-value threshold, i.e., 0.05.

| Cancer dataset | Distance | AUC ± SD<br>at P-value 5e-02 | Cancer dataset | Distance | AUC ± SD<br>at $L_p$ 2 |
| --- | --- | --- | --- | --- | --- |
| BRCA | $D_1$ | $0.75 \pm 0.11$ | BRCA | $D_1$ | $0.55 \pm 0.07$ |
| BRCA | $D_2$ | $0.62 \pm 0$ | BRCA | $D_2$ | $0.56 \pm 0.05$ |
| BRCA | $D_3$ | $0.68 \pm 0$ | BRCA | $D_3$ | $0.65 \pm 0.05$ |
| SKCM | $D_1$ | $0.78 \pm 0.26$ | SKCM | $D_1$ | $0.85 \pm \text{NA}$ |
| SKCM | $D_2$ | $0.78 \pm 0$ | SKCM | $D_2$ | $0.78 \pm \text{NA}$ |
| SKCM | $D_3$ | $0.78 \pm 0$ | SKCM | $D_3$ | $0.28 \pm \text{NA}$ |
| Basal-like vs HER2-enriched | $D_1$ | $0.90 \pm 0.08$ | Basal-like vs HER2-enriched | $D_1$ | $0.80 \pm \text{NA}$ |
| Basal-like vs HER2-enriched | $D_2$ | $0.78 \pm 0$ | Basal-like vs HER2-enriched | $D_2$ | $0.60 \pm \text{NA}$ |
| Basal-like vs HER2-enriched | $D_3$ | $0.28 \pm 0$ | Basal-like vs HER2-enriched | $D_3$ | $0.30 \pm \text{NA}$ |
| Basal-like vs Luminal-B | $D_1$ | $0.83 \pm 0.06$ | Basal-like vs Luminal-B | $D_1$ | $0.70 \pm 0.02$ |
| Basal-like vs Luminal-B | $D_2$ | $0.85 \pm 0$ | Basal-like vs Luminal-B | $D_2$ | $0.85 \pm 0.05$ |
| Basal-like vs Luminal-B | $D_3$ | $0.80 \pm 0$ | Basal-like vs Luminal-B | $D_3$ | $0.80 \pm 0.08$ |
| HER2-enriched vs Luminal-B | $D_1$ | $0.80 \pm 0.44$ | HER2-enriched vs Luminal-B | $D_1$ | $0.70 \pm \text{NA}$ |
| HER2-enriched vs Luminal-B | $D_2$ | $0.85 \pm 0$ | HER2-enriched vs Luminal-B | $D_2$ | $0.85 \pm \text{NA}$ |
| HER2-enriched vs Luminal-B | $D_3$ | $0.80 \pm 0$ | HER2-enriched vs Luminal-B | $D_3$ | $0.80 \pm \text{NA}$ |
| Basal vs like-Luminal-A | $D_1$ | $0.70 \pm 0.18$ | Basal vs like-Luminal-A | $D_1$ | $0.70 \pm 0$ |
| Basal vs like-Luminal-A | $D_2$ | $0.56 \pm 0.01$ | Basal vs like-Luminal-A | $D_2$ | $0.70 \pm 0.20$ |
| Basal vs like-Luminal-A | $D_3$ | $0.70 \pm 0$ | Basal vs like-Luminal-A | $D_3$ | $0.75 \pm 0.07$ |

**Table 3:** List of parameters in BIOMD0000000146 that can be mapped to proteins. Compared to the Biomodels version, the parameters were renumbered from 1 to 89, including the 11 conservation laws. The parameter sensitivities are provided as normalized average distances in 3 versions: *D*_1_ full distance, *D*_2_ distance along MAPKPP axis, *D*_3_ distance along AKTPiPP axis. *D*_1_ is computed using Eq. (16) (in main text), whereas *D*_2_ and *D*_3_ are computed based on Eq. (17) (in main text) with MAPKPP and AKTPiPP as target variables respectively. The tables in clockwise (starting with top-left) correspond to the *L_p_* norm values of 0.01, 0.1, 0.5 and 1 respectively.

| Par | Protein | $D_1$ | $D_2$ | $D_3$ | Par | Protein | $D_1$ | $D_2$ | $D_3$ | Par | Protein | $D_1$ | $D_2$ | $D_3$ | Par | Protein | $D_1$ | $D_2$ | $D_3$ |
| --- | --- | --- | --- | --- | --- | --- | --- | --- | --- | --- | --- | --- | --- | --- | --- | --- | --- | --- | --- |
| $k_{77}$ | DUSP | 2.071e-58 | 0.610 | 0 | $k_{77}$ | DUSP | 9.729e-07 | 0.610 | 0 | $k_{77}$ | DUSP | 0.047 | 0.610 | 0 | $k_{77}$ | DUSP | 0.240 | 0.610 | 0 |
| $k_{78}$ | PP2A | 4.816e-34 | 0.945 | 1.000 | $k_{78}$ | PP2A | 1.000e-03 | 0.945 | 1.000 | $k_{78}$ | PP2A | 0.372 | 0.945 | 1.000 | $k_{78}$ | PP2A | 0.693 | 0.945 | 1.000 |
| $k_{79}$ | AKT3 | 6.663e-52 | 0 | 0.916 | $k_{79}$ | AKT3 | 5.227e-06 | 0 | 0.916 | $k_{79}$ | AKT3 | 0.097 | 0 | 0.916 | $k_{79}$ | AKT3 | 0.306 | 0 | 0.916 |
| $k_{81}$ | PI3K | 3.213e-19 | 0 | 0.122 | $k_{81}$ | PI3K | 7.642e-03 | 0 | 0.122 | $k_{81}$ | PI3K | 0.298 | 0 | 0.122 | $k_{81}$ | PI3K | 0.505 | 0 | 0.122 |
| $k_{82}$ | MAPK | 1.100e-58 | 1.000 | 0 | $k_{82}$ | MAPK | 3.006e-07 | 1.000 | 0 | $k_{82}$ | MAPK | 0.027 | 1.000 | 0 | $k_{82}$ | MAPK | 0.145 | 1.000 | 0 |
| $k_{83}$ | MEK | 1.912e-39 | 0.972 | 0 | $k_{83}$ | MEK | 9.667e-05 | 0.972 | 0 | $k_{83}$ | MEK | 0.141 | 0.972 | 0 | $k_{83}$ | MEK | 0.356 | 0.972 | 0 |
| $k_{84}$ | RAF | 8.895e-35 | 0.387 | 0 | $k_{84}$ | RAF | 2.408e-04 | 0.387 | 0 | $k_{84}$ | RAF | 0.095 | 0.387 | 0 | $k_{84}$ | RAF | 0.223 | 0.387 | 0 |
| $k_{85}$ | RAS | 6.039e-73 | 0 | 0 | $k_{85}$ | RAS | 1.986e-08 | 0 | 0 | $k_{85}$ | RAS | 0.008 | 0 | 0 | $k_{85}$ | RAS | 0.075 | 0 | 0 |
| $k_{86}$ | SOS/GRB2 | 1.000 | 0.720 | 0 | $k_{86}$ | SOS/GRB2 | 1.000 | 0.720 | 0 | $k_{86}$ | SOS/GRB2 | 0.837 | 0.720 | 0 | $k_{86}$ | SOS/GRB2 | 0.888 | 0.720 | 0 |
| $k_{87}$ | SHC | 0.997 | 0.944 | 0 | $k_{87}$ | SHC | 9.398e-01 | 0.944 | 0 | $k_{87}$ | SHC | 1.000 | 0.944 | 0 | $k_{87}$ | SHC | 1.000 | 0.944 | 0 |
| $k_{89}$ | EGFR | 2.950e-16 | 0 | 0.250 | $k_{89}$ | EGFR | 7.439e-02 | 0 | 0.250 | $k_{89}$ | EGFR | 0.682 | 0 | 0.250 | $k_{89}$ | EGFR | 0.784 | 0 | 0.250 |

**Table 4:** Effect of P-value threshold on the AUC values. In each column, the median AUC values are reported by averaging over different *L_p_* norm values (0.01, 0.1, 0.5, 1) while keeping the P-value thresholds constant. SD represents the standard deviation. AUC values at NA entries could not be computed due to the absence of significant proteins at the particular P-value threshold.

| Cancer dataset | Distance | AUC ± SD<br>at P-value 5e-05 | AUC ± SD<br>at P-value 5e-04 | AUC ± SD<br>at P-value 5e-03 |
| --- | --- | --- | --- | --- |
| BRCA | $D_1$ | $0.65 \pm 0.05$ | $0.65 \pm 0.05$ | $0.66 \pm 0.09$ |
| BRCA | $D_2$ | $0.50 \pm 0$ | $0.50 \pm 0$ | $0.63 \pm 0$ |
| BRCA | $D_3$ | $0.65 \pm 0$ | $0.65 \pm 0$ | $0.55 \pm 0$ |
| SKCM | $D_1$ | NA | NA | NA |
| SKCM | $D_2$ | NA | NA | NA |
| SKCM | $D_3$ | NA | NA | NA |
| Basal-like vs HER2-enriched | $D_1$ | NA | NA | NA |
| Basal-like vs HER2-enriched | $D_2$ | NA | NA | NA |
| Basal-like vs HER2-enriched | $D_3$ | NA | NA | NA |
| Basal-like vs Luminal-B | $D_1$ | $0.80 \pm 0.04$ | $0.80 \pm 0.04$ | $0.80 \pm 0.01$ |
| Basal-like vs Luminal-B | $D_2$ | $0.85 \pm 0$ | $0.85 \pm 0$ | $0.85 \pm 0$ |
| Basal-like vs Luminal-B | $D_3$ | $0.80 \pm 0$ | $0.80 \pm 0$ | $0.80 \pm 0$ |
| HER2-enriched vs Luminal-B | $D_1$ | NA | NA | NA |
| HER2-enriched vs Luminal-B | $D_2$ | NA | NA | NA |
| HER2-enriched vs Luminal-B | $D_3$ | NA | NA | NA |
| Basal vs like-Luminal-A | $D_1$ | NA | NA | $0.80 \pm 0.04$ |
| Basal vs like-Luminal-A | $D_2$ | NA | NA | $0.85 \pm 0$ |
| Basal vs like-Luminal-A | $D_3$ | NA | NA | $0.80 \pm 0$ |

**Table 5:** Effect of *L_p_* norm on the AUC values. In each column, the median AUC values are reported by averaging over different P-value thresholds (5e-05, 5e-04, 0.005) while keeping the *L_p_* norm constant. SD represents the standard deviation. SD values at NA entries could not be computed due to the presence of significant proteins only at a single P-value threshold, i.e., 0.05.

| Cancer dataset | Distance | AUC ± SD<br>at $L_p$ 0.01 | AUC ± SD<br>at $L_p$ 0.1 | AUC ± SD<br>at $L_p$ 0.5 | AUC ± SD<br>at $L_p$ 1 |
| --- | --- | --- | --- | --- | --- |
| BRCA | $D_1$ | $0.70 \pm 0.03$ | $0.70 \pm 0.03$ | $0.65 \pm 0.04$ | $0.60 \pm 0.01$ |
| BRCA | $D_2$ | $0.56 \pm 0.07$ | $0.56 \pm 0.05$ | $0.56 \pm 0.05$ | $0.56 \pm 0.05$ |
| BRCA | $D_3$ | $0.65 \pm 0.05$ | $0.65 \pm 0.05$ | $0.65 \pm 0.05$ | $0.65 \pm 0.05$ |
| SKCM | $D_1$ | $0.35 \pm NA$ | $0.35 \pm NA$ | $0.78 \pm NA$ | $0.85 \pm NA$ |
| SKCM | $D_2$ | $0.78 \pm NA$ | $0.78 \pm NA$ | $0.78 \pm NA$ | $0.78 \pm NA$ |
| SKCM | $D_3$ | $0.28 \pm NA$ | $0.28 \pm NA$ | $0.28 \pm NA$ | $0.28 \pm NA$ |
| Basal-like vs HER2-enriched | $D_1$ | $1.00 \pm NA$ | $1.00 \pm NA$ | $0.90 \pm NA$ | $0.90 \pm NA$ |
| Basal-like vs HER2-enriched | $D_2$ | $0.60 \pm NA$ | $0.60 \pm NA$ | $0.60 \pm NA$ | $0.60 \pm NA$ |
| Basal-like vs HER2-enriched | $D_3$ | $0.30 \pm NA$ | $0.30 \pm NA$ | $0.30 \pm NA$ | $0.30 \pm NA$ |
| Basal-like vs Luminal-B | $D_1$ | $0.80 \pm 0.05$ | $0.80 \pm 0.05$ | $0.80 \pm 0.01$ | $0.80 \pm 0.01$ |
| Basal-like vs Luminal-B | $D_2$ | $0.85 \pm 0.05$ | $0.85 \pm 0.05$ | $0.85 \pm 0.05$ | $0.85 \pm 0.05$ |
| Basal-like vs Luminal-B | $D_3$ | $0.80 \pm 0.08$ | $0.80 \pm 0.08$ | $0.80 \pm 0.08$ | $0.80 \pm 0.08$ |
| HER2-enriched vs Luminal-B | $D_1$ | $0.80 \pm NA$ | $0.80 \pm NA$ | $0.80 \pm NA$ | $0.80 \pm NA$ |
| HER2-enriched vs Luminal-B | $D_2$ | $0.85 \pm NA$ | $0.85 \pm NA$ | $0.85 \pm NA$ | $0.85 \pm NA$ |
| HER2-enriched vs Luminal-B | $D_3$ | $0.80 \pm NA$ | $0.80 \pm NA$ | $0.80 \pm NA$ | $0.80 \pm NA$ |
| Basal vs like-Luminal-A | $D_1$ | $0.76 \pm 0.04$ | $0.76 \pm 0.04$ | $0.75 \pm 0.07$ | $0.75 \pm 0.07$ |
| Basal vs like-Luminal-A | $D_2$ | $0.70 \pm 0.20$ | $0.70 \pm 0.20$ | $0.70 \pm 0.20$ | $0.69 \pm 0.22$ |
| Basal vs like-Luminal-A | $D_3$ | $0.75 \pm 0.07$ | $0.75 \pm 0.07$ | $0.75 \pm 0.07$ | $0.75 \pm 0.07$ |

### 4.4 CPTAC Proteomics Data

CPTAC or Clinical Proteomic Tumor Analysis Consortium is another proteomic collaboration, containing TCGA patients. The CPTAC proteomic datasets relevant to the TCGA were downloaded from CPTAC data portal [1]. From all TCGA cancer datasets, only Breast, Ovarian and Colorectal cancers were available in CPTAC. Since we need relative values for our analysis, we used data derived from iTRAQ (isobaric Tags for Relative and Absolute Quantification) quantification method. More information about iTRAQ method and overall data analysis pipelines of CPTAC can be found in [2, 3]. The data consisted of log fold ratio of the samples (determined with respect to an internal reference).

#### 4.4.1 CPTAC Differential Protein Expression Analysis

We focused on CPTAC datasets with at least two groups of patient cohorts and the availability of iTRAQ data type. This resulted in a single dataset, namely, Breast cancer dataset with four subtypes (Luminal A, Luminal B, Basal-like and HER2-enriched subtypes). Using the linear modeling of the Limma package, we computed the differentially expressed proteins among the four subtypes in a pairwise manner. For each protein we calculated average log2 expression, moderated t-statistic, P-value, q-value or adjusted P-value (based on Benjamini and Hochberg correction method), log-odds that the protein is differentially expressed, and estimate of the log2-fold-change by applying empirical Bayes moderation of the standard errors towards a common value. More information can be found on Limma’s documentation and vignettes available at Bioconductor [4]. The significant proteins are determined based on various P-value thresholds, namely, 5e-05, 5e-04, 0.005, 0.05 and subsequently the significant model parameters. The adjusted P-values are reported in Table 1.

#### 4.4.2 Validating parameter sensitivity scores with CPTAC database

Out of 89 model parameters, we could map 11 parameters with the datasets. The AUC values were computed as per Section 4.3.3 and are reported in Table 2 and Supplementary Tables 4, 5. The distribution of AUC values are shown in Supplementary Fig. 6, 7, 5.

## 5 Discussion and conclusion

Computing the sensitivity scores based on tropical geometry is a flexible method for predicting sensitive parameters and tipping points of biochemical networks. The comparison between these predictions and differential proteomics data is summarized in the Table 2 and Supplementary Tables 4, 5.

We have used two types of sensitivity scores: an *n*-dimensional (nD) score based on the *L_p_* distance between sets of tropical equilibration solutions and a 1-dimensional (1D) score based on distances between projections of these sets on a target variable. Among these scores, the one based on the distance *D*_3_, the 1D score targeting AKT-PI-PP, has the worst performance in terms of the AUC of the ROC curve. The nD score and the 1D score targeting MAPK-PP lead to reasonable agreement between predicted sensitive parameters and proteomic data.

We measure the strength of overlap of our scores with the protein measurement data using AUC values. An AUC above 0.5 means that our method performs better than random guessing. We achieved better AUC values for many datasets, e.g., Luminal A, Luminal B, Basal-like and HER2-enriched subtypes from CPTAC database as compared to BRCA and SKCM datasets from TCPA database. For associating our scores with protein measurement data, two important parameters need to be pre-defined, namely *L_p_* norm for distance computation and the P-value threshold to assess protein significance. Furthermore, it has been suggested that in high dimensional setting, the proper choice of the *L_p_* norm is important [7], which was also reflected in our histograms (cf. Fig. 2). Therefore, we systematically varied the P-value thresholds along with the *L_p_* norm values resulting in different combinations and associated AUC values (cf. Fig. 4, 5, 6, 7). For a fixed *L_p_* we average over the different P-value thresholds and report the median AUC values along with the standard deviations. In the presence of significant proteins at only a single P-value threshold, the standard deviations are denoted as NA (cf. Table 2 and Supplementary Table 4). Likewise, for fixed P-value thresholds, we average over different *L_p_* norms. In the absence of significant proteins at a given threshold, AUC computation fails and is denoted by NA (cf. Table 2 and Supplementary Tables 5). We found that for most datasets a low value of *p* in the *L_p_* resulted in better AUC values, except in SKCM. The empirical findings suggest that a proper selection of the *L_p_* norm is important. Also, it was found that the choice of the P-value threshold is relevant.

**Fig. 4:**
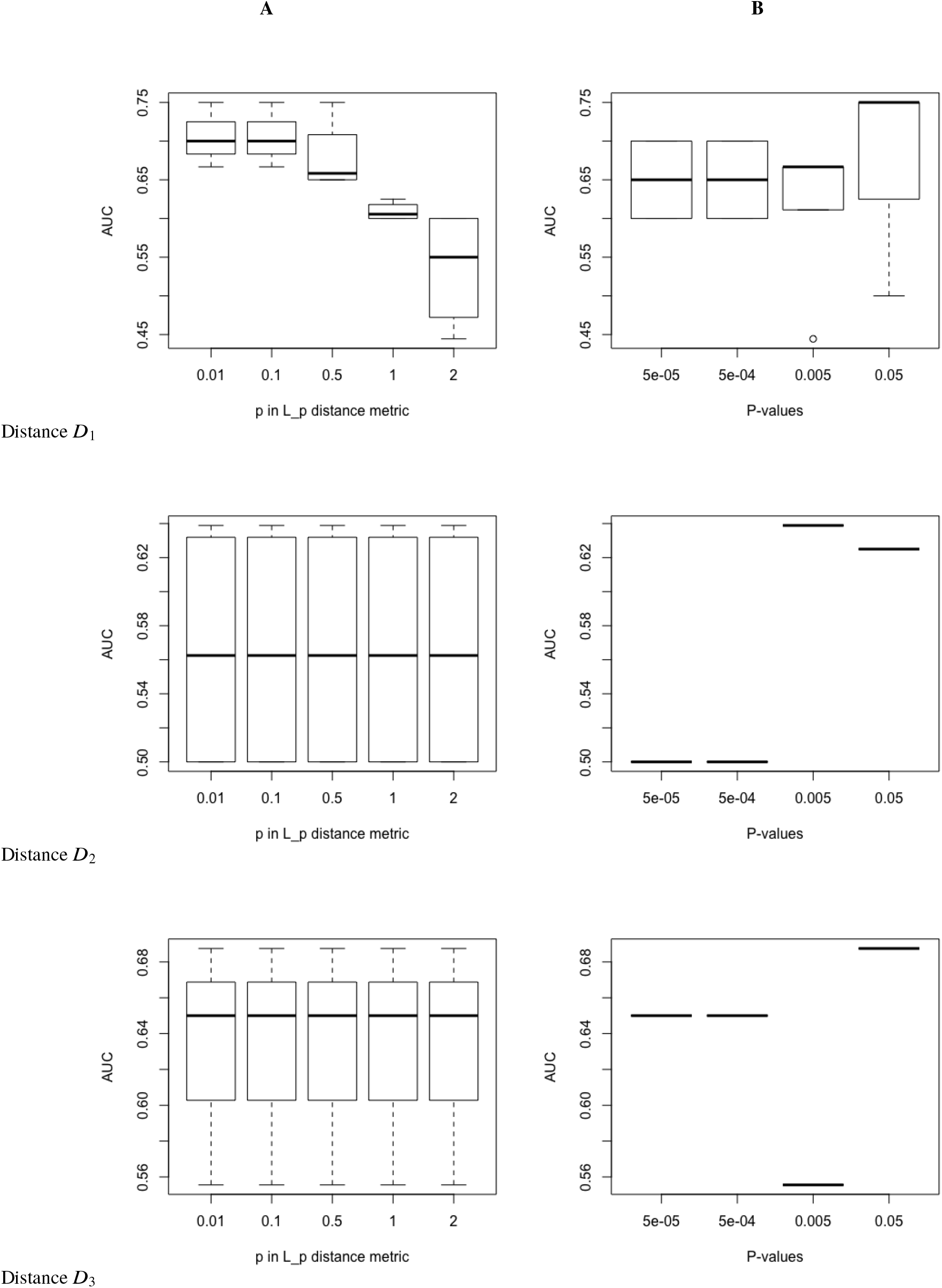
Distribution of AUC values for BRCA dataset obtained from TCPA database. Column **(A)**: The AUCs were computed for different P-value thresholds (cf. Section 4.3.2 and 4.3.3 in main text) for fixed values of *L_p_* norm and vice-versa in the column **(B)**. The AUCs are computed at distance measures *D*_1_, *D*_2_, *D*_3_ (cf. Supplementary Table 3 and Fig. 2 in main text).

**Fig. 5:**
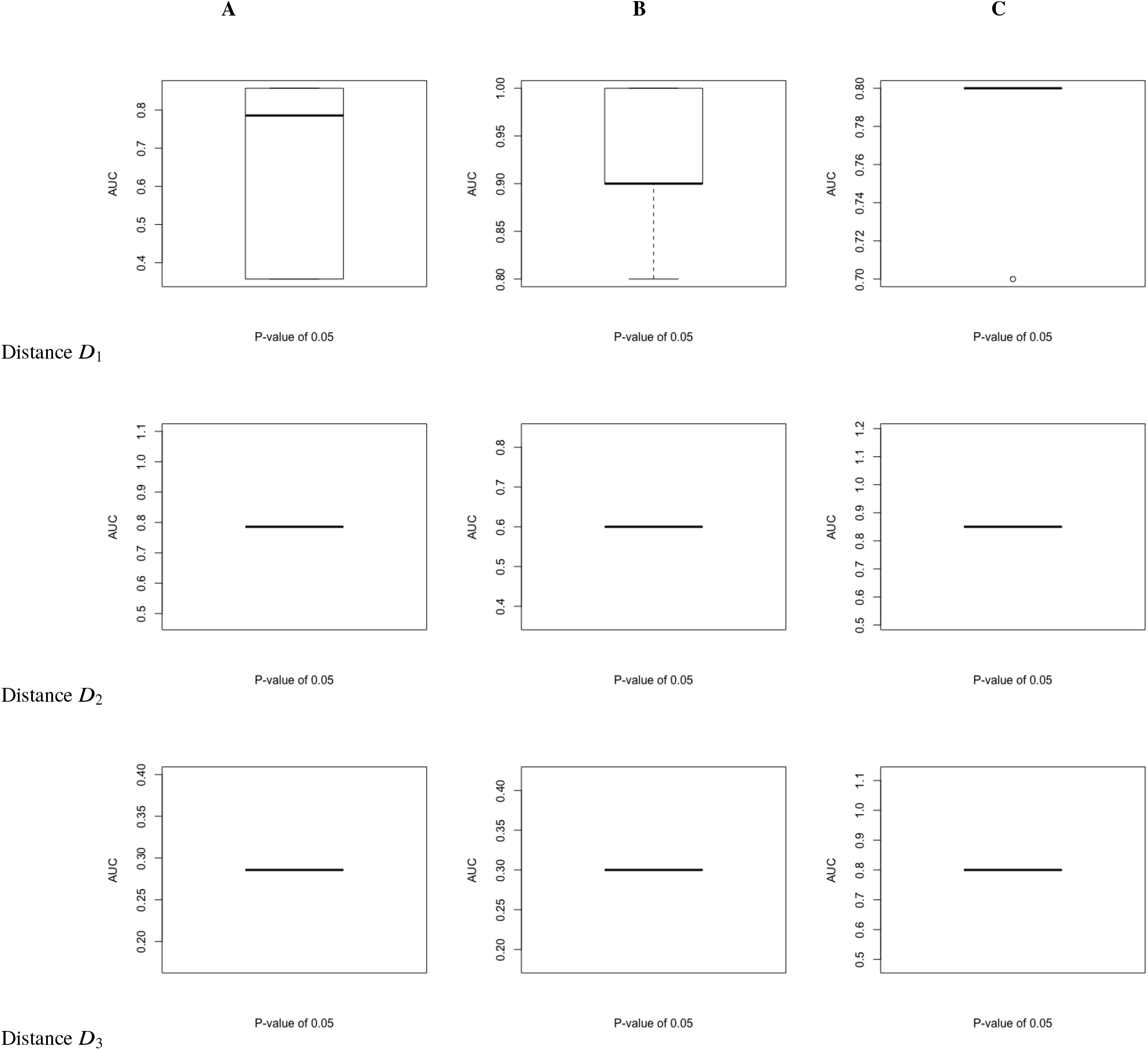
Figure shows distribution of AUC values from different databases: column **(A)**: for SKCM dataset obtained from TCPA database; row **(B)**: for Basal-like vs HER2-enriched breast cancer subtypes in CPTAC database and row **(C)** for HER2-enriched vs Luminal B breast cancer subtypes in CPTAC database. The AUCs were computed for different values of *L_p_* norm (0.01, 0.1, 0.5, 1, 2) for a fixed P-value threshold (cf. Sections 4.3.2 and 4.3.3, 4.4.1 and 4.4.2, 4.4.1 and 4.4.2 in main text for the corresponding dataset). The AUCs are computed at distance measures *D*_1_, *D*_2_, *D*_3_ (cf. Supplementary Table 3 and Fig. 2 in main text). We detected no significant proteins below the P-value threshold of 0.05.

**Fig. 6:**
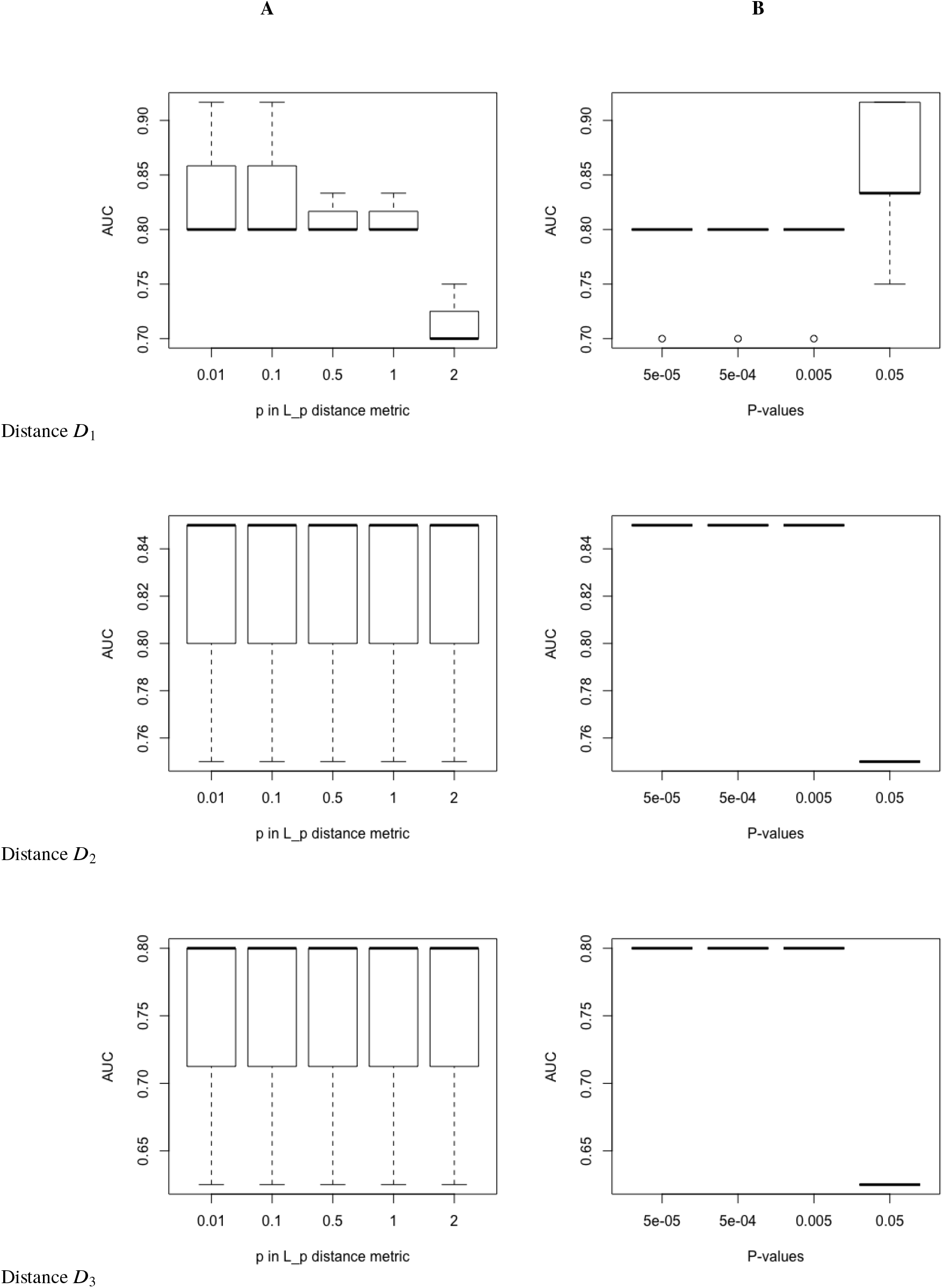
Distribution of AUC values for Basal-like vs Luminal-B breast cancer subtypes in CPTAC database. In the left, the AUCs were computed for different P-value thresholds for fixed values of *L_p_* norm and vice-versa in the right figure. The AUCs are computed at distance measures *D*_1_, *D*_2_, *D*_3_ (cf. Supplementary Table 3 and Fig. 2 in main text).

**Fig. 7:**
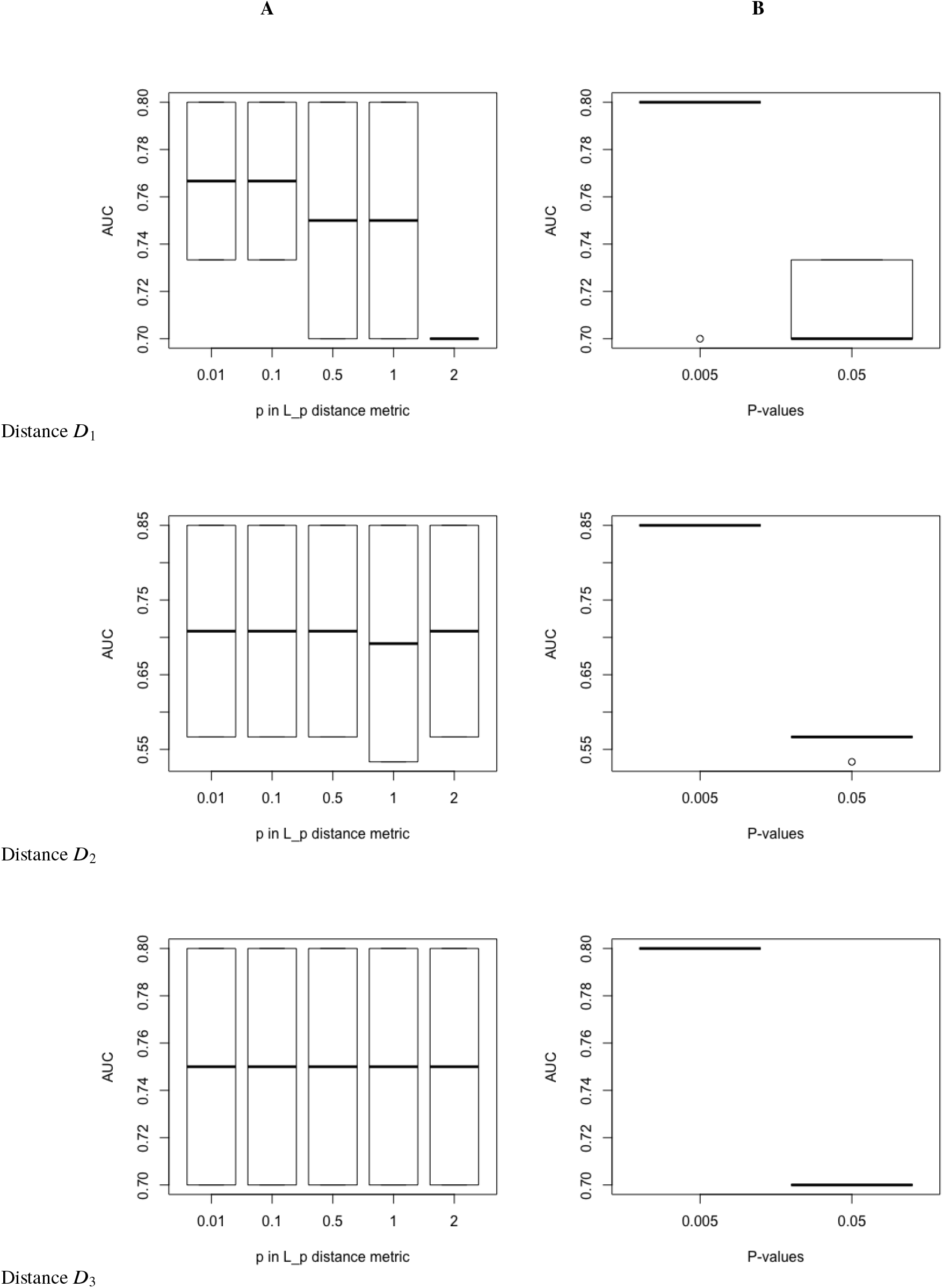
Distribution of AUC values for Basal-like vs Luminal-A breast cancer subtypes in CPTAC database. Column **(A)**: AUCs were computed for different P-value thresholds for fixed values of *L_p_* norm and vice-versa in the column **(B)** (cf. Section 4.4.1 and 4.4.2 in main text). We detected no significant proteins below the P-value threshold of 0.005. The AUCs are computed at distance measures *D*_1_, *D*_2_, *D*_3_ (cf. Supplementary Table 3 and Fig. 2 in main text).

The nD score with *p* = 0.01 for the *L_p_* norm identifies SOS/GRB2 and SHC as sensitive parameters. EGFR is not signaled as sensitive parameter, except for much larger *p* = 2, which means that EGFR total quantity is not limiting in this model. Extra sensitive parameters such as MAPK, MEK, PP2A and DUSP are detected by the 1D score when MAPK-PP is the target. PP2A, AKT3 are sensitive parameters when the target is AKT-PI-PP. The findings for 1D scores are compatible with standard sensitivity analysis of the same model. For instance, [18] found that the most influential parameters on MAPK-PP are, in decreasing order, MAPK, SHC, MEK, DUSP, and the most influential parameters on AKT-PI-PP are PP2A, AKT, PI3K, EGFR. This result is nearly identical to our Table 2 (columns *D*_2_ and *D*_3_; our *D*_2_ score classifies also PP2A among the parameters influential on MAPK-PP).

Altogether, the findings for nD and 1D scores are compatible with the properties of the studied model. The sensitive parameter PP2A is responsible for the behavior of the model as a single pathway (MAPK) or as two pathways in crosstalk [16]. The negative loop produced by the repression of MAPK by the PI3K/Akt pathway (via Raf) reduces the model sensitivity with respect to the total EGFR, but does not affect the model sensitivity on parameters downstream of Raf, such as MEK, MAPK and DUSP. The system of coupled MAPK and PI3K/Akt signaling networks has a number of other negative loops, not represented in this model. These loops further increase the robustness of this system and reduce the effectiveness of targeted therapies.

Contrary to standard sensitivity analysis methods, tropical sensitivity analysis has the choice to tune the contrast between sensible parameters by the choice of the *L_p_* norm, which can lead to well separated sensitivities where standard methods find a continuous distribution of sensitivities. Also, tropical sensitivity analysis does not use simulation of trajectories as standard sensitivity analysis does and is thus protected against errors coming from incomplete sampling of the trajectory set. Furthermore, as tropical analysis depends on orders of magnitude of the parameters, its results are expected to be valid for wide domains of model parameters. Finally, tropical sensitivity analysis can be also applied to categorical variables. For instance, one can compare patients with distinct types of mutations.

Tropical sensitivity analysis provides robust information on sensitive model parameters that are candidates for biologically important drivers. The extension of our approach to perturbation of multi parameter sets (e.g., combinatorial perturbations) remains a topic of future work. Furthermore, we aim to use drug-target databases for further in-silico validations of our sensitivity scores. We also aim to efficiently extrapolate our findings to gene expression databases which are more readily available than protein measurement databases. We also intend to study the effect of drug inhibitors in targeted therapy. In a tropical geometry setting, this can be studied by supplementing the model with variables representing the drugs and parameters representing the affinities of the drugs to the targets. Finally, a future topic is the study of the rich geometric structure of the perturbed metastable regimes resulting from the tropical sensitivity analysis. For instance, the geometrical description of the perturbed pathways could be used to characterize the pathway remodeling induced by the perturbation. We expect that refinement of these techniques will lead to a better understanding of tumor vulnerabilities that can be used to optimize targeted therapy.

## Acknowlegements

This work was supported by the ANR/DFG grant ANR-17-CE40-0036 (project SYMBIONT) and CompSE profile area, RWTH Aachen University. We thank R. Larive and D. Santamaria for their critical reading of the manuscript and for useful discussions.

## References

1. CPTAC data portal. https://proteomics.cancer.gov/data-portal.

2. CPTAC data portal. https://cptac-data-portal.georgetown.edu/cptac/documents/CDAP_Results_Overview_rev_09152014.pdf.

3. CPTAC data portal. https://cptac-data-portal.georgetown.edu/cptac/documents/CDAP_ProteinReports_description_20160503.pdf.

4. Limma page on Bioconductor. https://bioconductor.org/packages/release/bioc/html/limma.html.

5. TCGA data portal. https://cancergenome.nih.gov/abouttcga.

6. TCPA data portal. http://bioinformatics.mdanderson.org/main/TCPA:Overview.

7. Charu C. Aggarwal, Alexander Hinneburg, and Daniel A. Keim. On the surprising behavior of distance metrics in high dimensional space. In Jan Van den Bussche and Victor Vianu, editors, Database Theory — ICDT 2001, pages 420–434, Berlin, Heidelberg, 2001. Springer Berlin Heidelberg.

8. Nir Atias, Sorin Istrail, and Roded Sharan. Pathway-based analysis of genomic variation data. Current opinion in genetics & development, 23(6):622–626, 2013.

9. Yoav Benjamini and Yosef Hochberg. Controlling the false discovery rate: A practical and powerful approach to multiple testing. Journal of the Royal Statistical Society. Series B (Methodological), 57(1):289–300, 1995.

10. Francis S Collins and Harold Varmus. A new initiative on precision medicine. New England Journal of Medicine, 372(9):793–795, 2015.

11. Robin D Dowell, Owen Ryan, An Jansen, Doris Cheung, Sudeep Agarwala, Timothy Danford, Douglas A Bernstein, P Alexander Rolfe, Lawrence E Heisler, Brian Chin, et al. Genotype to phenotype: a complex problem. Science, 328(5977):469–469, 2010.

12. Hajk-Georg Drost. philentropy: Similarity and Distance Quantification Between Probability Functions, 2018. R package version 0.1.0.

13. Raphaela Fritsche-Guenther, Franziska Witzel, Anja Sieber, Ricarda Herr, Nadine Schmidt, Sandra Braun, Tilman Brummer, Christine Sers, and Nils Blüthgen. Strong negative feedback from Erk to Raf confers robustness to MAPK signalling. Molecular systems biology, 7(1):489, 2011.

14. Albeter Goldbeter and DE Koshland. Ultrasensitivity in biochemical systems controlled by covalent modification. interplay between zero-order and multistep effects. Journal of Biological Chemistry, 259(23):14441–14447, 1984.

15. Luca Grieco, Laurence Calzone, Isabelle Bernard-Pierrot, François Radvanyi, Brigitte Kahn-Perles, and Denis Thieffry. Integrative modelling of the influence of MAPK network on cancer cell fate decision. PLoS computational biology, 9(10):e1003286, 2013.

16. Mariko Hatakeyama, Shuhei Kimura, Takashi Naka, Takuji Kawasaki, Noriko Yumoto, Mio Ichikawa, Jae-Hoon Kim, Kazuki Saito, Mihoro Saeki, Mikako Shirouzu, et al. A computational model on the modulation of mitogen-activated protein kinase (MAPK) and Akt pathways in heregulin-induced ErbB signalling. Biochemical Journal, 373(Pt 2):451, 2003.

17. Myles Hollander and Douglas A Wolfe. Nonparametric statistical methods. 1999.

18. Dawei Hu and Jian-Min Yuan. Time-dependent sensitivity analysis of biological networks: coupled MAPK and PI3K signal transduction pathways. The journal of physical chemistry A, 110(16):5361–5370, 2006.

19. Bertram Klinger, Anja Sieber, Raphaela Fritsche-Guenther, Franziska Witzel, Leanne Berry, Dirk Schumacher, Yibing Yan, Pawel Durek, Mark Merchant, Reinhold Schäfer, et al. Network quantification of EGFR signaling unveils potential for targeted combination therapy. Molecular systems biology, 9(1):673, 2013.

20. Steven M LaValle. Planning algorithms. Cambridge university press, 2006.

21. Nicolas Le Novere, Benjamin Bornstein, Alexander Broicher, Melanie Courtot, Marco Donizelli, Harish Dharuri, Lu Li, Herbert Sauro, Maria Schilstra, Bruce Shapiro, Jacky L. Snoep, and Michael Hucka. BioModels database: a free, centralized database of curated, published, quantitative kinetic models of biochemical and cellular systems. Nucleic Acids Research, 34(suppl 1):D689–D691, 2006.

22. Jun Li, Yiling Lu, Rehan Akbani, Zhenlin Ju, Paul L Roebuck, Wenbin Liu, Ji-Yeon Yang, Bradley M Broom, Roeland GW Verhaak, David W Kane, et al. TCPA: a resource for cancer functional proteomics data. Nature methods, 10(11):1046, 2013.

23. Christoph Lüders. PtCut: Calculate tropical equilibrations and prevarieties. http://www.wrogn.com/ptcut/.

24. V. Noel, D. Grigoriev, S. Vakulenko, and O. Radulescu. Tropical geometries and dynamics of biochemical networks. Application to hybrid cell cycle models. Electronic Notes in Theoretical Computer Science, 284:75–91, 2012.

25. Nancy A. Obuchowski. Receiver operating characteristic curves and their use in radiology. Radiology, 229(1):3–8, 2003. PMID: 14519861.

26. Ovidiu Radulescu, Satya Swarup Samal, Aurélien Naldi, Dima Grigoriev, and Andreas Weber. Symbolic dynamics of biochemical pathways as finite states machines. In International Conference on Computational Methods in Systems Biology, pages 104–120. Springer, 2015.

27. Radulescu, O., Vakulenko, S., and Grigoriev, D. Model reduction of biochemical reactions networks by tropical analysis methods. Math. Model. Nat. Phenom., 10(3):124–138, 2015.

28. Aurélien Rizk, Gregory Batt, François Fages, and Sylvain Soliman. A general computational method for robustness analysis with applications to synthetic gene networks. Bioinformatics, 25(12):i169–i178, 2009.

29. Xavier Robin, Natacha Turck, Alexandre Hainard, Natalia Tiberti, Frédérique Lisacek, Jean-Charles Sanchez, and Markus Müller. pROC: an open-source package for R and S+ to analyze and compare ROC curves. BMC Bioinformatics, 12:77, 2011.

30. Suzanne L Rutherford. From genotype to phenotype: buffering mechanisms and the storage of genetic information. Bioessays, 22(12):1095–1105, 2000.

31. Satya Swarup Samal, Dima Grigoriev, Holger Fröhlich, and Ovidiu Radulescu. Analysis of reaction network systems using tropical geometry. In Vladimir P. Gerdt, Wolfram Koepf, Werner M. Seiler, and Evgenii V. Vorozhtsov, editors, Computer Algebra in Scientific Computing - 17th International Workshop (CASC 2015), volume 9301 of Lecture Notes in Computer Science, pages 422–437, Aachen, Germany, September 2015. Springer.

32. Satya Swarup Samal, Dima Grigoriev, Holger Fröhlich, Andreas Weber, and Ovidiu Radulescu. A geometric method for model reduction of biochemical networks with polynomial rate functions. Bulletin of mathematical biology, 77(12):2180–2211, 2015.

33. Satya Swarup Samal, Aurélien Naldi, Dima Grigoriev, Andreas Weber, Nathalie Théret, and Ovidiu Radulescu. Geometric analysis of pathways dynamics: application to versatility of TGF-βreceptors. Biosystems, 149:3–14, 2016.

34. Martin L Sos, Stefanie Fischer, Roland Ullrich, Martin Peifer, Johannes M Heuckmann, Mirjam Koker, Stefanie Heynck, Isabel Stückrath, Jonathan Weiss, Florian Fischer, et al. Identifying genotype-dependent efficacy of single and combined PI3K-and MAPK-pathway inhibition in cancer. Proceedings of the National Academy of Sciences, 106(43):18351–18356, 2009.

35. Richard Strohman. Maneuvering in the complex path from genotype to phenotype. Science, 296(5568):701–703, 2002.

36. Jeongah Yoon and Thomas S Deisboeck. Investigating differential dynamics of the MAPK signaling cascade using a multi-parametric global sensitivity analysis. PloS one, 4(2):e4560, 2009.

37. Zhike Zi. Sensitivity analysis approaches applied to systems biology models. IET systems biology, 5(6):336–346, 2011.

